# Clustering of mRNA-Seq data for detection of alternative splicing patterns

**DOI:** 10.1101/021733

**Authors:** Marla Johnson, Elizabeth Purdom

## Abstract

Current sequencing of mRNA can provide estimates of the levels of individual isoforms within the cell, where isoforms are the different distinct mRNA products or proteins created by a gene. It remains to adapt many standard statistical methods commonly used for analyzing gene expression levels to take advantage of this additional information. One novel question is whether we can find groupings or clusters of samples that are distinguished not by their gene expression but by their isoform usage. Such clusters in tumors, for example, could be the result of shared disruption to the splicing system that creates the different isoforms. We propose a novel approach to clustering mRNA-Seq data that identifies clusters of samples with common isoform usage. We show via simulation that our methods are more sensitive to finding clusters of similar alternative splicing patterns than standard clustering techniques applied directly to the estimates of isoform levels. We further demonstrate that clustering on isoform usage is more accurate than clustering directly on isoform levels by examining real data that contains a technical artifact that resulted in different batches having different isoform usage patterns. Clustering, mRNA-Seq, Alternative splicing

## 1 Introduction

Clustering techniques are widely used in gene expression studies, ranging from visualization of the data (in the form of heatmaps) to more formal detection of underlying subpopulations. Cancer studies, for example, often rely on clustering to detect subtypes of tumors based on the expression patterns of genes (see for example Perou *and others* (2000); Sorlie *and others* (2001)). These subtypes can have important clinical properties, creating an important link between the biological mechanisms of tumor cells with the phenotypes observed in tumor patients.

There are obviously many methods for clustering of gene expression, and classical techniques such as hierarchical clustering and k-means or mediods are frequently used. Traditionally, gene expression studies were based on microarray studies, but now even large studies often measure gene expression levels by sequencing of mRNA (Hammerman *and others*, 2012; The Cancer Genome Atlas Research Network, 2013).

Measuring gene expression using mRNA-sequencing, rather than microarray technologies, introduces some added statistical complexity to a clustering analysis. The most obvious is that RNA-sequencing often results in estimates that are the count of the number of sequences from each gene (depending on how the estimates are determined). These estimates are integer valued and non-negative, and do not follow the standard log-normal assumptions of microarray data. In the context of estimating differential expression between pre-defined groups, there has been much work in accounting for the statistical properties of these estimates (Robinson and Smyth, 2007; Anders and Huber, 2010; Zhou *and others*, 2011; Yang *and others*, 2012; Wu *and others*, 2013; Yu *and others*, 2013; Leng *and others*, 2013). In the context of clustering, there has been some work in utilizing appropriate model-based methods for count data, including the Poisson and Negative binomial distributions (Witten, 2011; Si *and others*, 2014), though it is still quite common to log-transform the data and apply standard clustering techniques.

Another novel aspect of using mRNA-sequencing for measuring expression levels is that sequencing provides a wealth of information about alternative splicing within the cell. Alternative splicing is the process by which a single gene codes for multiple mRNA products (or *isoforms*) by the process of including or excluding portions of the DNA of a gene (examples of which include exon skipping, intron retention, or alternative 5’ and 3’ splice sites). Different isoforms within a gene often have different functions which are location and development specific. Some proteins even have isoforms that have antagonisti functions; for example, the gene VEGF has one isoform that is used by cancer cells to encourage new vasculature near tumors and also an anti-angiogenic form that inhibits tumor growth (Qiu *and others*, 2009).

With mRNA-sequencing, expression levels of individual isoforms can be estimated, rather than just the cumulative level from the entire gene. Despite the possible importance of individual isoforms, clustering on mRNA-Seq data often still relies on gene estimates – i.e. the total amount of mRNA, summed over all isoforms of the gene – rather than individual isoform-level expression data. If there are not many genes with isoform differences in the samples under consideration, then a clustering based upon thousands of genes is unlikely to change regardless of whether clustering is based on isoform or gene expression levels.

However, in some settings, there might be large scale changes in splicing. For example, in the setting of cancer studies, there is evidence of abnormalities in tumors that might have a wide spread effect on the splicing across many genes. For example, dysregulation in the machinery of the cell that controls splicing (the spliceosome) could have splicing impacts on many isoforms; significantly mutated genes that are members of the spliceosome have been found in several tumors (Yoshida *and others*, 2011; Je *and others*, 2013; Makishima *and others*, 2012). An example is the gene SF3B1 which encodes for a subunit of the spliceosome, and has been found to have mutations that lead to aberrant splicing in around 15-20% and 10% of uveal melanoma and chronic lymphocytic leukemia tumors, respectively (Furney *and others*, 2013; Gentien *and others*, 2014; Quesada *and others*, 2012). However, mutation is just one in which function of a gene can be disregulated in tumors; it is well known that similar phenotypes in tumors can be the result of abnormalities other than mutations. This suggests that unsupervised clustering techniques, which do not rely on identifying the source of the abnormality, could provide greater ability to detect dysregulated splicing on tumors. This is particularly true given that significant mutations in the spliceosome have been found at much lower prevelance for other tumors, such as gene U2AF1 which is found mutated at a prevalence of only around 5% in acute myleoid lymphoma or lung adenocarcinoma (The Cancer Genome Atlas Research Network, 2013; Cancer Genome Atlas Research Network, 2014).

There are multiple ways to incorporate the information provided by individual isoform estimates into clustering. The most obvious is to cluster based on isoform estimates, rather than gene estimates. Isoform estimates are similar in structure to that of gene estimates and therefore such clustering could make use of similar existing procedures. Our strategy here, however, is on evaluating the *relative* isoform usage within a gene: a measure of the tendency of a gene to prefer one isoform over another. This leads to a data structure where the isoforms are grouped by gene. This is a different data structure than the traditional *n × p* data matrix, for which the corresponding clustering methods are not appropriate.

While the focus of this paper is entirely on the biological application of clustering isoform usage, it can be helpful to posit this problem into a more general framework. This question of clustering relative isoform usage can be stated more generally as the problem of clustering data when the features are known to have a predefined grouping assignment. Using this terminology, gene estimates are then a summary statistic (the sum) of the features in the group. Then gene expression clustering is clustering of a summary statistic of the group members, while isoform clustering is clustering of the individual features ignoring the group membership.

We propose a clustering strategy that uses the group structure in a flexible manner. We assume that within each group there is a natural notion of distance between samples based on the features in that group; for isoform usage, as we discuss below, we use the distance between the estimates of proportional isoform usage within the gene. We then create distance metrics separately for each group (or gene), and then aggregate the distances across groups and apply standard clustering techniques on the aggregate distance. This creates a flexible clustering strategy that allows the feature information to contribute to the clustering only in the context of its relationship to other features in the group. Our focus in what follows will be entirely on the specific example of clustering of isoform usage, but it is useful to note that other choices of distances with a group could capture different kinds of relationships for clustering.

We evaluate this clustering strategy on simulated isoform expression data and show that it has improved performance in detecting isoform usage changes, as compared to clustering on the individual features (isoform clustering) or to clustering on the summary statistic (gene clustering). We also demonstrate our clustering technique on a mRNA-Seq dataset that has a clear technical artifact which appears to have affected the isoform usage; using this as a gold-standard, we show that our clustering strategy detects the underlying clustering more accurately than the other two strategies.

### 1.1 Brief Biological background

We give a brief biological background to alternative splicing so as to make clear the biological terms we use.

#### Alternative Splicing

The DNA that contains the genetic code for proteins must be first copied or transcribed into a free floating messenger RNA (mRNA) that is then transported to the ribosomes and converted into a protein. In eukaryotic cells, the process of copying the DNA into mRNA has itself stages so that a copy of the DNA is first made (called pre-mRNA) which is then changed into the final mRNA that makes the protein. In many eukaryotes, the changes induced on the pre-mRNA includes selectively cutting out portions of the pre-mRNA so that the final (mature) mRNA transcript is no longer an exact copy of the code found in the DNA of the cell. The process of cutting out portions of the pre-mRNA is called *splicing*. In many complex organisms, including human and many common model organisms like fruit flies and mice, there is additional complexity so that there are multiple ways in which the same pre-mRNA can be spliced; in addition, there are also multiple pre-mRNA that can be made from the same gene, depending on which of several starting points and ending points are used in copying DNA into pre-mRNA. The end result is that a single gene representing a stretch of code on the DNA can result in many different transcripts, or *isoforms* of the gene.

#### Sequencing of mRNA

Previously, large-scale quantification of the amount of mRNA in a cell (also called *mRNA expression*) was via microarray technology; expression measurements from microarrays quantified all mRNA from a gene without distinguishing between different isoforms of the gene. The rapid expanse in sequencing technologies has allowed for direct sequencing of mRNA in order to determine the amount of each unique mRNA in a cell, and therefore opens up the ability to quantify not just the cumulative amount of expression from a gene region, but also that of individual isoforms within a gene. However, it is important to note that current commonly used sequencing technologies still do not allow for the entire mRNA to be sequenced; instead the mRNA must be cut into smaller fragments that are then sequenced. This means that estimates of the amount of individual isoforms are not the simple result of counting how many sequences came from particular isoform, but must be indirectly estimated via deconvolution methods (Denoeud *and others*, 2008; Jiang and Wong, 2009; Trapnell *and others*, 2010; Richard *and others*, 2010; Salzman *and others*, 2010; Katz *and others*, 2010). Such methods provide with an estimate of the underlying rate of transcription of each isoform, which can then be translated into “estimated counts” so that they are on the same scale *as if* one could have uniquely identified the sequences to isoforms and simply counted them.

### 2 Methods

We are interested in the effect of clustering based on *isoform usage*, by which we mean the relative percentage within each gene that an isoform is used. Using our terminology from the introduction, our features are individual isoforms which are grouped a priori into genes. Specifically, for a single sample, each group consists of a vector of features, one for each isoform of the gene. We can think of the gene as defining a grouping of the isoforms, and we want the comparisons between isoforms to be within the gene group so that comparisons of isoforms across genes is mediated by their relative behavior within the gene. In what follows, we will focus our method and terminology solely on clustering based on differential isoform usage, though the concepts could be generalized.

More formally, for each sample *i* and gene *j* we observed not a single value, but a vector of values *p*_*ij*_*∈ R*^*K*_*j*_^, where *K*_*j*_ is the number of features or isoforms in gene *j*. We note that *K*_*j*_ varies across genes, and we only consider genes with multiple isoforms so that *K*_*j*_*>* 1. In the data we considered, *K*_*j*_ ranged from 2 isoforms up to instances of more than 30 expressed isoforms.

Let the estimates of isoform expression for individual *i* in gene *j* be denoted as *x*_*ij*1_, *…, x_ijK_j* and let 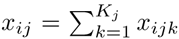 be the total expression in gene *j*, i.e. the estimate of gene expression. Then our estimate of isoform usage within gene *j* for sample *i* is given by the relative proportion of each isoform within the gene, 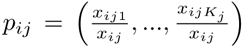 We note that for our clustering method we could consider any vector *p*_*ij*_ that is a function of our isoform information, including for example the original, untransformed vector of isoform expression values, but our focus is on the relative expression within the gene so we focus on the proportion. Gonzàlez-Porta *and others* (2012) similarly conver isoform estimates to proportions per gene to quantify variability in splicing and identify those genes with varying splicing ratios.

As a result, our data of isoform usage is clearly not a simple *n × p* matrix assumed by many clustering programs. Of course, many clustering techniques need simply a distance matrix between the *n* samples for clustering, but it is also not obvious how to create a distance matrix between the samples when each of the vectors *p*_*ij*_ lies in an entirely different dimensional space.

However, for a single gene (or group), there are numerous distances defined between vectors that lie on the simplex. Therefore, we take the strategy of calculating per gene distances *d*_*j*_(*p*_*ij*_, *p*_*i'j*_) and aggreagate the different *d*_*j*_(*p*_*ij*_, *p*_*i'j*_) across genes. There are obviously also many choices for aggregating the *J* distance matrices, and a straightforward strategy is to define a aggregate distance based on a (weighted) average of the different distance matrices,

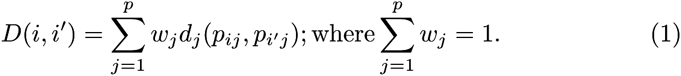

In the results that follow, we ultimately weighted each feature equally so that *D*(*i, i'*) was a simple average of the distances calculated per gene – that is,.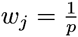

This same strategy is suggested by the objective functions of many clustering routines, where the objective function to be minimized can be written as an average over individual terms that involve only the per feature distance matrices. For example, for K-means clustering the standard objective function is to determine clusters *C*_*k*_ that minimize the within cluster distances; for standard Euclidean distance, which can be written as *d*(*i, i'*) = Σ_*j*_*d*_*j*_(*i, i'*), this implies (Witten and Tibshirani, 2010),

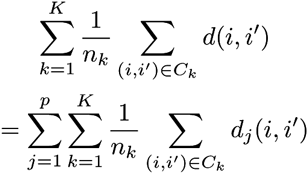

For other clustering techniques, the resulting aggregate distance object, *D*(*i, i'*) can be converted via multidimensional scaling (MDS) into a two-dimensional data matrix preserving the distances, which could also be used as input into clustering algorithms.

By adopting this framework, we now only need to define a distance for relative isoform usage *per feature*. This means that we simply need a relevant distance between vectors of proportions, for which there are many options (see Deza and Deza, 2013, for a review); Gonzàlez-Porta *and others* (2012) use Hellinger’s distance in their analysis of isoform proportions. We consider some of the most popular: *χ*^2^ distance, euclidean distance, Jeffrey’s divergence, and Hellinger’s distance; we also consider a distance based on the log-likelihood (Berninger *and* others, 2008; Witten, 2011) to account for the difference in variability due to different counts (see Supplemental Text, Section 1 for more details).

#### Weighting of features

In our implementation, we ultimately weighted each feature equally, but as we note above, different weights for each group, or gene, can be used rather than a simple average of the individual group distances. As only a small fraction of features differ between clusters, identifying and weighting these genes based on the data may lead to improved clustering and interpretable weights; Witten and Tibshirani (2010) proposed a sparse clustering technique that tries to find a sparse set of such weights by putting a *L*_1_ penalty on the weights and iteratively finding the best weights. When we generalizing the sparse methods of (Witten and Tibshirani, 2010) to find weights automatically in our clustering, this added an enormous amount of variability in our clustering results and therefore did not perform well even in simple situations.

#### Relationship to Kernel Methods

Our strategy is also related to kernel strategies. Specifically, because of the relationship between distances and kernels, we can see this approach as defining a separate kernel for each gene, and then combining the kernels via (weighted) averaging. Methods for combining multiple kernels have been proposed and how to choose functionals that combine multiple kernels is termed the multiple kernel learning problem (see Gönen and Alpaydin, 2011, for a review), and weighted combinations like we describe are common, Many of these methods have computational difficult for the large numbers of features we have here. Zeng and Cheung (2010) propose creating a sparse multiple kernel for each feature for unsupervised clustering, with the similar goal as Witten and Tibshirani (2010) of finding sparse weights for the linear combinations of the kernels for each feature.

### 3 Results

#### 3.1 Simulation Study

We evaluate via simulation the performance of clustering on gene expression, isoform expression, or proportion levels. To test the ability of the gene, isoform, and proportion clustering algorithms, we jointly simulated gene counts and corresponding isoform proportions under the following scenarios for how a gene could show clustering,

1. The gene expression counts different between groups, while the proportion levels are constant across groups. In this case, the isoform expression will also vary but will be consistent with changes in gene expression. We would expect the gene and isoform clustering methods to distinguish these groups but the proportion clustering method to not differentiate them.
2. The gene expression is constant across samples, while the proportion levels vary between groups. In this case, the isoform expression levels will also vary due to the changes in isoform relative frequency. We expect the proportion and isoform clustering methods to distinguish these groups but the gene clustering method to not distinguish them.
3. Both the gene expression and proportion levels vary across groups. In this case, isoform expression will vary due to changes in both gene expression and isoform relative frequency. We expect all methods (gene, isoform, and proportion), to distinguish clustering patterns, though not necessarily the same clusters depending if the proportion and gene clusters do not overlap.

##### 3.1.1 Description of Simulation

We set up simulations of 5,000 genes across 135 samples, with the number of isoforms within each gene simulated from a Poisson(2) distribution, truncated to be strictly greater than 2; this was to roughly mimid the distribution seen in real mRNA Seq data. For each simulation, a percentage of the genes were given a set clustering pattern in their gene expression and/or isoform proportions depending on which of the above scenarios was being considered, while the remaining genes had both constant gene and isoform expression (no clustering signal). In order to see how sensitive the algorithm was, the percentage of genes with the pattern varied from 0.5%, 1%, 2%, 4%, 8%, and 10%. When both gene and proportion differences were seen in the same gene, we defined distinct clusters for gene expression from those of proportional isoform usage, where the proportional usage clusters were either nested within the gene expression clusters, or spaned across the gene clusters.

We also consider a more complicated scenario where different genes behave differently, with some showing clustering, as in Case 1 above, with only gene expression differences, and other showing both gene and alternative splicing differences, as in Case 3 from above. This is probably more realistic biologically, since there are likely to be many genes with gene expression differences without differences in isoform usage. To examine the effects of these conflicting signals, we simulated 5,000 genes where 25% of the genes had differential gene expression, while a varying number of genes (0.5%, 1%, 2%, 4%, 8%, and 10%) showed both differential gene expression and differential alternative splicing in their clustering, with the remaining genes simulated to have no clustering signal in their gene expression nor isoform usage. Again the gene expression clusters and the proportional usage clusters were distinct from each other, as described above.

To create the data, we simulated proportion vectors by randomly sampling isoform values and calculating their relative proportions. We then generated gene counts by similarly randomly sampling from isoform values and calculating their sums. The proportion vectors and gene counts were multiplied to return the final simulated isoform counts. These two separate stages (rather than just directly sampling isoform counts and getting both proportion and gene measures from those) allows us to separately control the gene level and the proportion level which permited more complicated interactions between the two. The isoform values in both cases were generated from a negative binomial and the clusters were created by changing the parameters of the negative binomial between clusters (whether between proportion or gene clusters); the parameters were based on means and dispersion parameters estimated from TCGA count data.

For further details regarding the simulations, see the Supplemental Text, Section 2.

##### 3.1.2 Results

The results of the simulations show that the proportion clustering reliably finds proportion clusters even when the pattern represents a low percentage of genes in the data (around 2% of genes, Figure 1a). Moreover, the proportion clustering finds the pattern even with different complicated backgrounds of other clustering signals based on gene expression differences (Figure 1b). These are both important characteristics, since we expect that differences in alternative splicing will affect a comparatively small numbers of genes and differences in overall gene expression will dominate.

**Figure 1:**
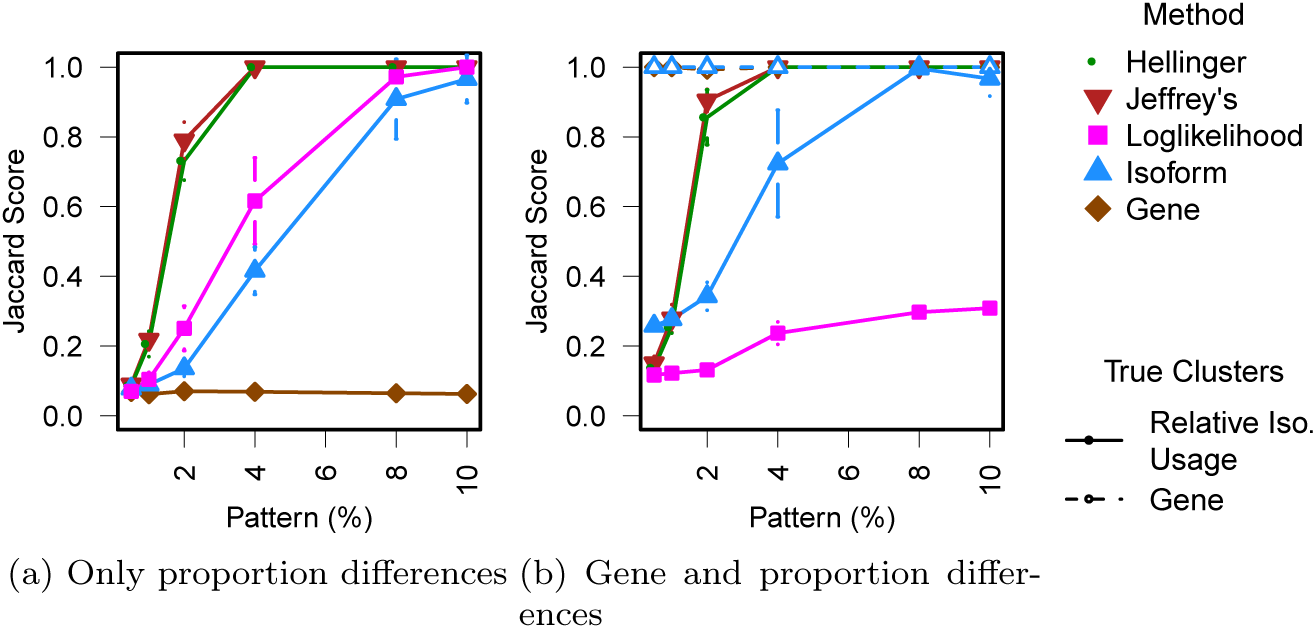
Jaccard Scores from Simulation Study. Plotted are the mean Jaccard Scores over 1,000 simulations (y-axis) against the percent of genes with the clustering pattern varies (x-axis). The Jaccard score is the proportion of pairs of samples correctly co-clustered, with one indicating perfect detection of the clusters. Different lines correspond to different clustering methods, indicated by the legend. ‘Hellinger’, ‘Jeffrey’s’ and ‘Loglikelihood’ refer to different choices for the distance taken in clustering on the proportions; ‘Isoform’ and ‘Gene’ refer to standard clustering on the isoform or gene expression estimates directly. Vertical bars stretch to ± 2 standard errors. (a) shows the results of the clustering when the only clustering signal comes from changes in proportional isoform usage within the gene (as expected, there is no signal from gene clustering). (b) shows the results of clustering when there are a complicated mix of clustering signals; specifically 25% of the genes show different gene expressions according to 3 gene expression clusters, and a varying percentage of different genes (as indicated in the x-axis) show both the different gene expression patterns according to the 3 gene expression clusters as well as different proportional isoform usage according to 6 non-nested proportional usage clusters (see Supplemental Text, Section 2 for a more complete description). The dotted lines show the Jaccard scores when trying to find the gene clusters, while solid lines show those from trying to find the proportional usage clusters. Shown here are the results from K-mediods as the clustering method; results from other scenarios and from hierarchical clustering are shown in Supplemental Figures S2-S5.

In compare differences between choices in the proportion clustering, we see that hierarchical clustering tends to do slightly better than K-mediods clustering (Supplementary Figure S3). Similarly, Jeffrey’s divergence performs slightly better than Hellinger distance. The distance based on log-likelihood performs poorly, often not finding the clusters at all. Chi-squared distance and Euclidean distances perform similarly to Jeffrey’s divergence and Hellinger, respectively (Supplementary Figure S1), and for simplicity are not shown on most of the figures from the simulations.

We next consider the performance of proportion clustering with that of clustering isoform expressions directly. Clustering on the isoform levels directly captures groups differences in overall gene expressions quite readily. This is clearly true when the only signal in the data comes from differences in overall gene expression, where clustering on isoforms has equivalent performance to that of gene (Supplemental Figure S2). And even when there are competing signals, corresponding to differences in relative proportional isoform use within the gene, the isoform generally finds the clusters with differing gene expressions well (Figure 1b and Supplemental Figures S4 and S5), except for the case when the proportional usage clusters clash with that of the gene clusters.

In contrast, clustering on isoform expression does not do as good of a job of finding clusters that differ in proportional isoform usage. When the only clustering differences are based on proportional isoform usage, clustering on the isoforms does not find the clusters until the genes with the signal are a relatively larger percentage of the data (Figure 1a). This is in contrast with the performance of proportion clustering, where the groups are found even when a small percentage of the genes show the signal.

In the more complicated settings we examine, where gene expression and proportional usage clusters co-exist, the isoform clustering performs slightly better, though still not as good as the clustering on the proportional isoform usage. The improvement is likely due to the fact that the proportional usage clusters share some clustering signal with the gene expression clusters, so that the isoform’s detection of the gene expression clustering improves its performance. Clustering on the isoform expression patternss only starts to have high concordence with the true proportional usage clusters when about 7-10% of the genes show that pattern – a much higher required percentage than that of the proportion clustering which finds the pattern reliably even when only 2% of the genes show the pattern.

These result indicate that proportion clustering has the potential to be more sensitive to finding clustering based on this type of alternative splicing, and particularly when there are a mix of gene and alternative splicing signals as would be expected in true biological data.

### 3.2 Clustering of LAML data

In general, it is difficult to compare clustering methods on real data absent knowledge of true groupings in the sample. Finding data with such groupings is quite difficult for our method, since it is difficult to know a priori that there are differences in relative isoform usage. However, in analyzing mRNA-Seq data of Acute Myeloid Leukemia (LAML) tumors, sequenced as part of the The Cancer Genome Atlas (TCGA) project (The Cancer Genome Atlas Research Network, 2013), we discovered an unreported batch effect in the data. Further exploration of this batch effect suggests that the effect on the data was likely to have resulted in different relative levels of isoform abundance within genes, as we explain below. This gives us a gold-standard to which we can compare the efficiency of our our clustering methods. We find that clustering on the relative isoform usage (i.e. proportions) in this dataset gives a practically exact correspondence to this batch effect and performs much better in finding this batch effect than clustering based on isoform levels.

#### 3.2.1 Implementation of the Clustering

We normalize the LAML samples by TMM normalization (Robinson and Oshlack, 2010), and we perform an initial filtering of the data to remove extremely lowly expressed isoforms. Using the isoform counts, we create the three different types of features: gene counts, isoform counts, and isoform proportions. The number of isoforms and genes provided is still quite large – 28,014 expressed isoforms and 12,218 expressed genes. We then apply a variance filter to reduce the size of the datasets to the 5,000 most variable isoforms or genes (which is a common practice in clustering of gene expression datasets). In the case of the isoform proportions, we filter by calculating the distance matrix for all features (i.e. genes) and chose the 5,000 genes with the largest summed distance matrix.

To provide greater stability in our clustering and to not be influenced by outlying samples, we performed consensus clustering (Monti *and others*, 2003) on top of our clustering routines. Briefly, this involves repeated subsampling of the entire dataset and enumerating how frequently individuals were clustered together in a consensus matrix. After performing 1000 subsamples, the consensus matrix is clustered to achieve our final clustering.

See Supplementary Text, Section 3 for further details regarding the implementation of the clustering methods on the LAML data.

#### 3.2.2 Results

In Figure 2 we compare the clustering assignments from clustering on the the three different features (gene, isoform, and proportion) in order to demonstrate the differences in how the samples are clustered. In Figure 2 we have also superimposed the “plate” from which the sample originated, which signifies the batch in which the samples were sent to be sequenced. We can see that all of the methods’ cluster assignments have a significant relationship with plate; plate 734 clusters together in all of the methods. Observing such an effect not terribly surprising, since effects due to different batches are quite common in large projects such as these, though the accompanying AML paper from which this data is drawn explicitly states that this data did not show a batch effect due to plate (The Cancer Genome Atlas Research Network, 2013).

**Figure 2:**
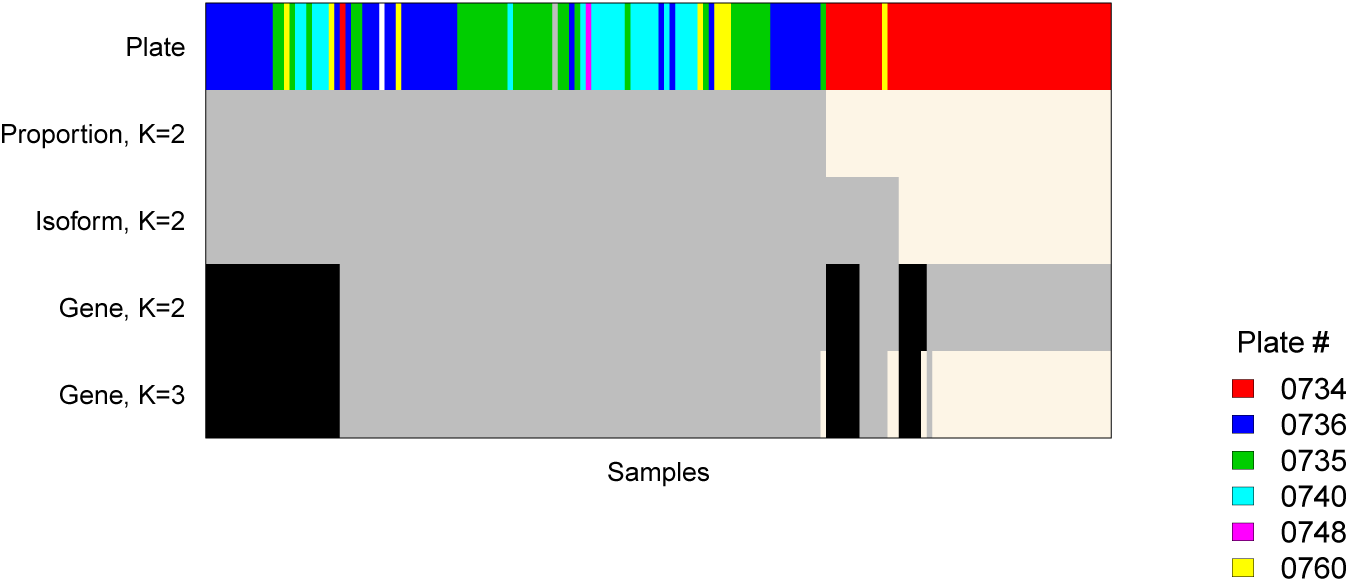
Comparison of Clustering Assignment. Each column corresponds to a sample and each row to a clustering method. The cluster assignments of the three different clusterings (isoform, gene and proportion) are shown for *K* = 2 groups by coloring the sample according to its clustering in the above plot. We also show *K* = 3 for gene clustering, since this is the point at which the gene clustering starts to have clusterings corresponding to the plate. The samples have been ordered to highlight the similarity between the clusterings. The top row shows the plate assignment of each sample, showing an almost perfect correspondence with proportion clustering. This clustering is based on concensus clustering of hierarchical clustering.

In comparing the performance of the methods in detecting the plate effect, it is clear that proportion clustering is almost perfectly detecting the plate effect, which is clearly its primary signal. The other two methods do not have as clear a correspondence with plate, regardless of what *K* is used or the method of clustering (kmeans or hierarchical). This is true for K-means clustering as well (Supplemental Figure S6), though the accordence of proportion clustering with the plate effect is not quite as good as with hierarchical clustering. Removal of the plate effect with the batch correction tool ComBat (Johnson *and others*, 2007), results in quite different clusterings, and the resulting clusterings have a much closer relationship with the other clinical data on the samples. This suggests that this effect is a technical artifact and has an adverse affect on the clustering results (the correlation between clinical variables and the clustering results shown in the accompanying AML paper closely match our results *after* we removed the plate effect, perhaps suggesting that they did perform some batch removal with their data, contrary to their statement).

While it is encouraging to find strong concordence with an known grouping of the observations, it is unclear whether this demonstrates that the clustering of proportions is doing better at finding clusters of differential isoform usage. In particular, it is not clear why the plate effect would have anything to do with isoform usage within the gene. However, when we examine the plate effect more closely, we see that the differences in these plates can be clearly seen by looking at statistics that quantify the 5’ to 3’ bias of the mRNA-Seq data. The 5’ to 3’ bias refers to the fact that the technical steps in sequencing the data has the effect that the 3’ end of a transcript (i.e. the terminal end of the transcript) is more likely to be captured and sequenced than the 5’ end (the beginning) of a transcript, creating a bias in the amount of expression detected in different parts of the gene. Isoforms often differ in their starting and ending exons, and therefore different relative coverage of the beginning or end of a gene due to technical artifacts can mean that isoforms will get assigned different relative expression levels.

In Figure 3 we show a plot of the relative proportion of the mRNA-Seq sequences that came from the beginning of the transcript (5’ end) versus the end of the transcript (3’ end), as calculated by RSeQC (Wang *and others*, 2012). It is clear from this plot, that plate 734 has greater relative coverage of the beginning of the gene compared to the the other plates; Figure 3 shows short genes (0-1000 base pairs) but the effect can be seen in a range of gene lengths (Supplemental Figure S7), though it appears to perhaps be an effect seen on a fixed number of base pairs at beginning of the gene, since its seen in smaller and smaller percentiles of the beginning of the gene. Some differences can also be seen in the 3’ end of the genes, though not as consistently (again see Supplemental Figure S7).

**Figure 3:**
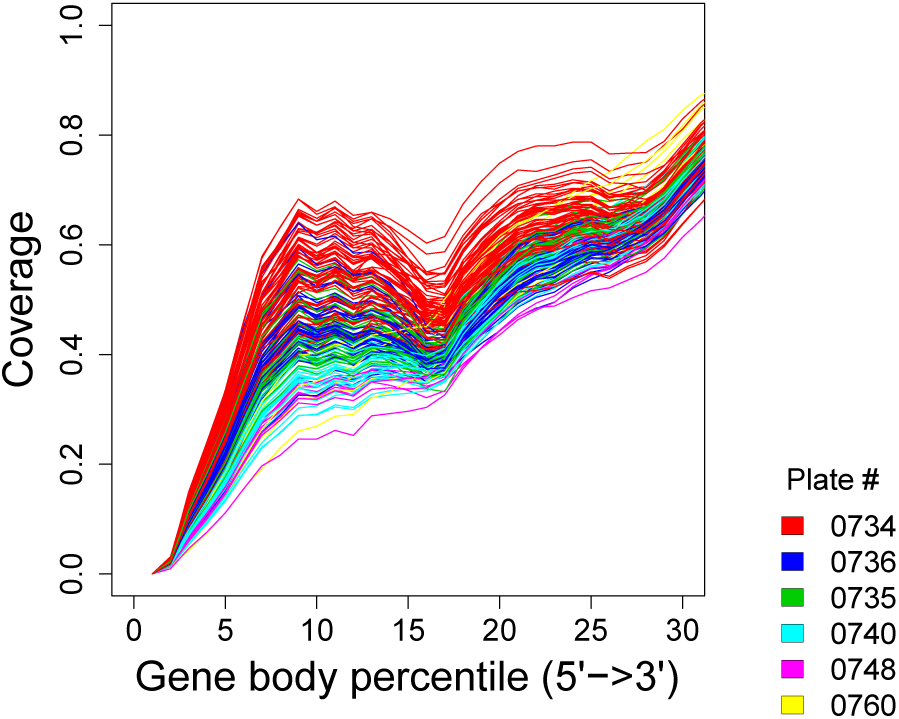
5’ to 3’ bias differs by plate. Here we show the results of RSeQC (Wang *and others*, 2012) calculation of the average coverage of the mRNA-Seq data; shown here are the results of the calculation for 318 housekeeping genes that have only a single isoform and have total length in the range of 0-1000bp. This calculation divides the gene into equally spaced bins and calculates the number of sequences falling in the bin, relative to the overall number sequences assigned to the gene region. The x-axis shows the percentile of the gene body that the bin falls in (referenced from the beginning, or 5’ end, of the gene). This plot shows a closeup of the results at the 5’ start of the gene; the plot showing the rest of the gene as well as plots showing genes of longer lengths are shown in Supplemental Figure S7.

Therefore, it is plausible that the 5’/3’ differences in the plates have created different relative expression of the isoforms, which the proportion clustering captures more accurately than the other methods.

## 4 Discussion

We have presented a method for clustering on isoform relative proportions so as to find shared patterns in alternative splicing. We demonstrate on simulated data that clustering on the proportions can more accurately detect changes in isoform usage within a gene than just clustering on isoform expression levels, even in the absense of any conflicting signal from gene expression. We further apply the method to a mRNA-Seq dataset which we demonstrate has a clear batch effect, and we see that clustering on relative proportions clusters the data by this batch effect, unlike the clustering from isoform or gene, which both imperfectly find the batch effect. Because the batch effect appears to create technical artifacts in the data that would influence isoform relative expression, this suggests that clustering on isoform relative proportions is more sensitive to correctly clustering the samples by differences in isoform usage.

Our method can be stated as a general strategy for clustering on features that have natural grouping structure, in this case where isoforms are grouped into genes. Our technique uses this grouping structure to define the relevant measure for comparing the features (or isoforms), and in this way focus the clustering on differences of features relative to the groups.

## 5 Supplementary Materials

The reader is referred to the on-line Supplementary Materials for supplementary text and figures.

### Funding

This work was supported by a Helman Faculty Grant, National Institutes of Health grant U24 CA143799 and National Science Foundation grant DMS- 1026441.

## Acknowledgements

The results published here are in part based upon data generated by The Cancer Genome Atlas pilot project established by the NCI and NHGRI. Information about TCGA and the investigators and institutions who constitute the TCGA research network can be found at http://cancergenome.nih.gov/.

## References

Anders, Simon and Huber, Wolfgang. (2010). Differential expression analysis for sequence count data. Genome Biology 11, R106.

Berninger, Philipp, Gaidatzis, Dimos, van Nimwegen, Erik and Za-volan, Mihaela. (2008). Computational analysis of small RNA cloning data. Methods 44(1), 13–21. MicroRNAs: Part B.

Cancer Genome Atlas Research Network. (2014, July). Comprehensive molecular profiling of lung adenocarcinoma. Nature 511(7511), 543–550.

Denoeud, France, Aury, Jean-Marc, Silva, Corinne Da, Noel, Ben-jamin, Rogier, Odile, Delledonne, Massimo, Morgante, Michele, Valle, Giorgio, Wincker, Patrick, Scarpelli, Claude, Jaillon, Olivier *and others*. (2008, Jan). Annotating genomes with massive-scale RNA sequencing. Genome Biol 9(12), R175.

Deza, Michel Marie and Deza, Elena. (2013, October). Encyclopedia of Distances, 2 edition. Springer.

Furney, Simon J, Pedersen, Malin, Gentien, David, Dumont, Amaury G, Rapinat, Audrey, Desjardins, Laurence, Turajlic, Samra, Piperno-Neumann, Sophie, de la Grange, Pierre, Roman-Roman, Sergio, Stern, Marc-Henri *and others*. (2013, October). SF3B1 mutations are associated with alternative splicing in uveal melanoma. Cancer discovery 3(10), 1122–1129.

Gentien, D, Kosmider, O, Nguyen-Khac, F, Albaud, B, Rapinat, A, Dumont, A G, Damm, F, Popova, T, Marais, R, Fontenay, M, Roman-Roman, S, Bernard, O A and others. (2014, January). A common alternative splicing signature is associated with SF3B1 mutations in malignancies from different cell lineages. Leukemia 28, 1355–1357.

Gönen, Mehmet and Alpaydin, Ethem. (2011). Multiple kernel learning algorithms. Journal of Machine Learning Research 12, 2211–2268.

Gonzàlez-Porta, Mar, Calvo, Miquel, Sammeth, Michael and Guigó, Roderic. (2012, March). Estimation of alternative splicing variability in human populations. Genome research 22(3), 528–538.

Hammerman, Peter S, Lawrence, Michael S, Voet, Douglas, Jing, Rui, Cibulskis, Kristian, Sivachenko, Andrey, Stojanov, Petar, Mckenna, Aaron, Lander, Eric S, Gabriel, Stacey, Getz, Gad, Sougnez, Carrie, Imielinski, Marcin, Helman, Elena, Hernan-dez, Bryan, Pho, Nam H, Meyerson, Matthew, Chu, Andy, Chun, Hye-Jung E, Mungall, Andrew J, Pleasance, Erin, Gordon Robertson, A, Sipahimalani, Payal, Stoll, Dominik, Balasun-daram, Miruna, Birol, Inanc, Butterfield, Yaron S N, Chuah, Eric, Coope, Robin J N, Corbett, Richard, Dhalla, Noreen, Guin, Ranabir, He, An, Hirst, Carrie, Hirst, Martin, Holt, Robert A, Lee, Darlene, Li, Haiyan I, Mayo, Michael, Moore, Richard A, Mungall, Karen, Ming Nip, Ka, Olshen, Adam, Schein, Jacqueline E, Slobodan, Jared R, Tam, Angela, Thiessen, Nina, Varhol, Richard, Zeng, Thomas, Zhao, Yongjun, Jones, Steven J M, Marra, Marco A, Saksena, Gordon, Cherniack, Andrew D, Schumacher, Stephen E, Tabak, Barbara, Carter, Scott L, Pho, Nam H, Nguyen, Huy, Onofrio, Robert C, Crenshaw, Andrew, Ardlie, Kristin, Beroukhim, Rameen, Winckler, Wendy, Ham-merman, Peter S, Getz, Gad, Meyerson, Matthew, Protopopov, Alexei, Zhang, Jianhua, Hadjipanayis, Angela, Lee, Semin, Xi, Ruibin, Yang, Lixing, Ren, Xiaojia, Zhang, Hailei, Shukla, Sa-chet, Chen, Peng-Chieh, Haseley, Psalm, Lee, Eunjung, Chin, Lynda, Park, Peter J, Kucherlapati, Raju, Socci, Nicholas D, Liang, Yupu, Schultz, Nikolaus, Borsu, Laetitia, Lash, Alex E, Viale, Agnes, Sander, Chris, Ladanyi, Marc, Todd Auman, J, Hoadley, Katherine A, Wilkerson, Matthew D, Shi, Yan, Liquori, Christina, Meng, Shaowu, Li, Ling, Turman, Yidi J, Topal, Michael D, Tan, Donghui, Waring, Scot, Buda, Elizabeth, Walsh, Jesse, Jones, Corbin D, Mieczkowski, Piotr A, Singh, Darshan, Wu, Junyuan, Gulabani, Anisha, Dolina, Peter, Bodenheimer, Tom, Hoyle, Alan P, Simons, Janae V, Soloway, Matthew G, Mose, Lisle E, Jefferys, Stuart R, Balu, Saianand, O’Connor, Brian D, Prins, Jan F, Liu, Jinze, Chiang, Derek Y, Neil Hayes, D, Perou, Charles M, Cope, Leslie, Danilova, Ludmila, Weisenberger, Daniel J, Maglinte, Dennis T, Pan, Fei, Van Den Berg, David J, Triche, Timothy Jr, Herman, James G, Baylin, Stephen B, Laird, Peter W, Getz, Gad, Noble, Michael, Voet, Doug, Saksena, Gordon, Gehlenborg, Nils, DiCara, Daniel, Zhang, Jinhua, Zhang, Hailei, Wu, Chang-Jiun, Yingchun Liu, Spring, Lawrence, Michael S, Zou, Lihua, Sivachenko, Andrey, Lin, Pei, Stojanov, Petar, Jing, Rui, Cho, Juok, Nazaire, Marc-Danie, Robinson, Jim, Thorvaldsdottir, Helga, Mesirov, Jill, Park, Peter J, Chin, Lynda, Schultz, Nikolaus, Sinha, Rileen, Ciriello, Giovanni, Cerami, Ethan, Gross, Benjamin, Jacobsen, Anders, Gao, Jianjiong, Arman Aksoy, B, Weinhold, Nils, Ramirez, Ricardo, Taylor, Barry S, Antipin, Yevgeniy, Reva, Boris, Shen, Ronglai, Mo, Qianxing, Seshan, Venkatraman, Paik, Paul K, Ladanyi, Marc, Sander, Chris, Akbani, Rehan, Zhang, Nianxiang, Broom, Bradley M, Casasent, Tod, Unruh, Anna, Wakefield, Chris, Craig Cason, R, Baggerly, Keith A, Weinstein, John N, Haussler, David, Benz, Christopher C, Stuart, Joshua M, Zhu, Jingchun, Szeto, Christopher, Scott, Gary K, Yau, Christina, Ng, Sam, Goldstein, Ted, Waltman, Peter, Sokolov, Artem, Ellrott, Kyle, Collisson, Eric A, Zerbino, Daniel, Wilks, Christopher, Ma, Singer, Craft, Brian, Wilkerson, Matthew D, Todd Auman, J, Hoadley, Katherine A, Du, Ying, Cabanski, Christopher, Walter, Vonn, Singh, Darshan, Wu, Junyuan, Gulabani, Anisha, Bodenheimer, Tom, Hoyle, Alan P, Simons, Janae V, Soloway, Matthew G, Mose, Lisle E, Jefferys, Stuart R, Balu, Saianand, Marron, J S, Liu, Yufeng, Wang, Kai, Liu, Jinze, Prins, Jan F, Neil Hayes, D, Perou, Charles M, Creighton, Chad J, Zhang, Yiqun, Travis, William D, Rekhtman, Natasha, Yi, Joanne, Aubry, Marie C, Cheney, Richard, Dacic, Sanja, Flieder, Douglas, Funkhouser, William, Illei, Peter, Myers, Jerome, Tsao, Ming-Sound, Penny, Robert, Mallery, David, Shelton, Troy, Hat-field, Martha, Morris, Scott, Yena, Peggy, Shelton, Candace, Sherman, Mark *and others*. (2012, September). Comprehensive genomic characterization of squamous cell lung cancers. Nature 489(7417), 519–525.

Je, Eun Mi, Yoo, Nam Jin, Kim, Yoo Jin, Kim, Myung Shin and Lee, Sug Hyung. (2013, July). Mutational analysis of splicing machinery genes SF3B1, U2AF1 and SRSF2 in myelodysplasia and other common tumors. International journal of cancer. Journal international du cancer 133(1), 260–265.

Jiang, Hui and Wong, Wing Hung. (2009, Apr). Statistical inferences for isoform expression in RNA-seq. Bioinformatics 25(8), 1026–32.

Johnson, W.E., Rabinovic, A. and Li, C. (2007). Adjusting batch effects in microarray expression data using empirical bayes methods. Biostatistics 8(1), 118–127.

Katz, Yarden, Wang, Eric T, Airoldi, Edoardo M and Burge, Christopher B. (2010, November). Analysis and design of RNA sequencing experiments for identifying isoform regulation. Nature Methods 7(12), 1009–1015.

Leng, Ning, Dawson, John A., Thomson, James A., Ruotti, Victor, Rissman, Anna I., Smits, Bart M. G., Haag, Jill D., Gould, Michael N., Stewart, Ron M. and Kendziorski, Christina. (2013, April). EBSeq: an empirical Bayes hierarchical model for inference in RNAseq experiments. Bioinformatics 29(8), 1035–1043.

Makishima, Hideki, Visconte, Valeria, Sakaguchi, Hirotoshi, Jankowska, Anna M, Abu Kar, Sarah, Jerez, Andres, Przychodzen, Bartlomiej, Bupathi, Manoj, Guinta, Kathryn, Afable, Manuel G, Sekeres, Mikkael A, Padgett, Richard A, Tiu, Ramon V *and others*. (2012, April). Mutations in the spliceosome machinery, a novel and ubiquitous pathway in leukemogenesis. Blood 119(14), 3203–3210.

Monti, S, Tamayo, P, Mesirov, J and Golub, T. (2003). Consensus clustering: a resampling-based method for class discovery and visualization of gene expression microarray data. Machine learning 52(1), 91–118.

Perou, C M, Sørlie, T, Eisen, M B, van de Rijn, M, Jeffrey, S S, Rees, C A, Pollack, J R, Ross, D T, Johnsen, H, Akslen, L A, Fluge, O, Pergamenschikov, A, Williams, C, Zhu, S X, Lønning, P E, Børresen-Dale, A L, Brown, P O *and others*. (2000, August). Molecular portraits of human breast tumours. Nature 406(6797), 747–752.

Qiu, Yan, Hoareau-Aveilla, Coralie, Oltean, Sebastian, Harper, Steven J and Bates, David O. (2009, December). The anti-angiogenic isoforms of VEGF in health and disease. Biochemical Society transactions 37(Pt 6), 1207–1213.

Quesada, Víctor, Conde, Laura, Villamor, Neus, Ordóñez, Gonzalo R, Jares, Pedro, Bassaganyas, Laia, Ramsay, Andrew J, Beà, Sílvia, Pinyol, Magda, Martínez-Trillos, Alejandra, LópezGuerra, Mónica, Colomer, Dolors, Navarro, Alba, Baumann, Tycho, Aymerich, Marta, Rozman, María, Delgado, Julio, Giné, Eva, Hernández, Jesús M, González-Díaz, Marcos, Puente, Di-ana A, Velasco, Gloria, Freije, José M P, Tubío, José M C, Royo, Romina, Gelpí, Josep L, Orozco, Modesto, Pisano, David G, Zamora, Jorge, Vázquez, Miguel, Valencia, Alfonso, Himmelbauer, Heinz, Bayés, Mónica, Heath, Simon, Gut, Marta, Gut, Ivo, Estivill, Xavier, López-Guillermo, Armando, Puente, Xose S, Campo, Elías *and others*. (2012, January). Exome sequencing identifies recurrent mutations of the splicing factor SF3B1 gene in chronic lymphocytic leukemia. Nature genetics 44(1), 47–52.

Richard, Hugues, Schulz, Marcel, Sultan, Marc, Nurnberger, Asja, Schrinner, Sabine, Balzereit, Daniela, Dagand, Emilie, Rasche, Axel, Lehrach, Hans, Vingron, Martin, Haas, Stefan *and others*. (2010, Jun). Prediction of alternative isoforms from exon expression levels in RNA-seq experiments. Nucleic Acids Research 38(10), e112.

Robinson, Mark and Oshlack, Alicia. (2010). A scaling normalization method for differential expression analysis of RNA-seq data. Genome Biology 11(3), R25–R25.

Robinson, Mark and Smyth, Gordon. (2007, Nov). Moderated statistical tests for assessing differences in tag abundance. Bioinformatics 23(21), 2881.

Salzman, Julia, Jiang, Hui and Wong, Wing Hung. (2010, March). Statistical modeling of RNA-SEQ data. Technical Report BIO-252, Division of Biostatistics, Stanford University, Palo Alto.

Si, Yaqing, Liu, Peng, Li, Pinghua and Brutnell, Thomas P. (2014, January). Model-based clustering for RNA-seq data. Bioinformatics (Oxford, England) 30(2), 197–205.

Sorlie, Therese, Perou, Charles M, Tibshirani, Robert, Aas, Turid, Geisler, Stephanie, Johnsen, Hilde, Hastie, Trevor, Eisen, Michael B, van de Rijn, Matt, Jeffrey, Stefanie S, Thorsen, Thor, Quist, Hanne, Matese, John C, Brown, Patrick O, Botstein, David, Lønning, Per Eystein and others. (2001). Gene Expression Patterns of Breast Carcinomas Distinguish Tumor Subclasses with Clinical Implications. Proceedings of the National Academy of Sciences of the United States of America 98(19), 10869–10874.

The Cancer Genome Atlas Research Network. (2013, May). Genomic and Epigenomic Landscapes of Adult De Novo Acute Myeloid Leukemia. The New England journal of medicine 368(22), 2059–2074.

Trapnell, C, Williams, B A, Pertea, G, Mortazavi, A, Kwan, G, van Baren, M J, Salzberg, S L, Wold, B J and Pachter, L. (2010, May). Transcript assembly and quantification by RNA-seq reveals unannotated transcripts and isoform switching during cell differentiation. Nature Biotechnology 28(5), 511.

Wang, L., Wang, S. and Li, W. (2012). Rseqc: quality control of rna-seq experiments. Bioinformatics 28, 2184–2185.

Witten, D M. (2011, December). Classification and clustering of sequencing data using a Poisson model. The Annals of Applied Statistics 5(4), 2493–2518.

Witten, D M and Tibshirani, R. (2010, June). A framework for feature selection in clustering. Journal of the American Statistical Association 105(490), 713–726.

Wu, Hao, Wang, Chi and Wu, Zhijin. (2013, April). A new shrinkage estimator for dispersion improves differential expression detection in RNA-seq data. Biostatistics 14(2), 232–243.

Yang, Xin, Todd, John A., Clayton, David and Wallace, Chris. (2012, November). Extra-binomial variation approach for analysis of pooled DNA sequencing data. Bioinformatics 28(22), 2898–2904.

Yoshida, Kenichi, Sanada, Masashi, Shiraishi, Yuichi, Nowak, Daniel, Nagata, Yasunobu, Yamamoto, Ryo, Sato, Yusuke, SatoOtsubo, Aiko, Kon, Ayana, Nagasaki, Masao, Chalkidis, George, Suzuki, Yutaka, Shiosaka, Masashi, Kawahata, Ryoichiro, Yamaguchi, Tomoyuki, Otsu, Makoto, Obara, Naoshi, SakataYanagimoto, Mamiko, Ishiyama, Ken, Mori, Hiraku, Nolte, Flo-rian, Hofmann, Wolf-Karsten, Miyawaki, Shuichi, Sugano, Sumio, Haferlach, Claudia, Koeffler, H Phillip, Shih, Lee-Yung, Haferlach, Torsten, Chiba, Shigeru, Nakauchi, Hiromitsu, Miyano, Satoru *and others*. (2011, October). Frequent pathway mutations of splicing machinery in myelodysplasia. Nature 478(7367), 64–69.

Yu, Danni, Huber, Wolfgang and Vitek, Olga. (2013, May). Shrinkage estimation of dispersion in Negative Binomial models for RNA-seq experiments with small sample size. Bioinformatics 29(10), 1275–1282.

Zeng, Hong and Cheung, Yiu-ming. (2010, November). Feature Selection and Kernel Learning for Local Learning Based Clustering. IEEE transactions on pattern analysis and machine intelligence 33(8), 1532–1547.

Zhou, Y H, Xia, K and Wright, F A. (2011, September). A powerful and flexible approach to the analysis of RNA sequence count data. Bioinformatics (Oxford, England) 27(19), 2672–2678.

